# Nanoscale architecture of malaria sporozoite formation

**DOI:** 10.64898/2026.11.24.768957

**Authors:** Esrah W. Du, Eleni Hughes, Jesse Aaron, W. Robert Shaw, Jeffrey D. Dvorin, Teng-Leong Chew, Flaminia Catteruccia

**Author notes:** **Corresponding author** Correspondence to Flaminia Catteruccia.

## Abstract

*Plasmodium falciparum* parasites rapidly form thousands of infectious daughter cells, called sporozoites, within single oocysts during their development in the mosquito midgut. How parasites accurately compartmentalize these daughter cells remains poorly understood, mostly due to the exceedingly large oocyst volume (circa 113,000 µm^3^ at the peak of sporozoite segmentation) and the substantial number of cells (5,000-10,000 sporozoites) hampering detailed ultrastructural studies^1^. Here, we use a pre-commercial focused ion beam-scanning electron microscopy platform and custom-trained deep learning models to reconstruct at nanometer resolution the overarching cellular ultrastructure of two oocysts during mid- and late sporozoite formation^2,3^. We unveil a highly synchronous segmentation process occurring with a striking degree of accuracy and reconstruct faithful stepwise interactions between key organelles in the different growth phases that likely ensure the fidelity of organellar segregation. We discover that sporozoites release membrane-bound cytoplasmic droplets, possibly to refine cell shape in preparation for their motility that drives transmission to humans. Contrary to previous assumptions, in both stages we conclusively visualize a single, interconnected sporoblast which we reason may help precisely coordinate signaling events that drive daughter cell formation. Our study provides foundational insight at unprecedented resolution into the ultrastructural features and changes characterizing the segmentation of *P. falciparum* sporozoites, conceivably the deadliest unicellular organisms in human history.

## Main

A defining feature of the *Plasmodium* life cycle that makes it such a prolific pathogen is periods of explosive population expansion resulting in the formation of multiple invasive daughter cells^4^. Rapid increases in biomass happen at three distinct points in parasite development: in hepatocytes during exoerythrocytic schizogony (up to 90,000 daughter liver merozoites)^5^, in blood stage schizogony (16-36 daughter merozoites)^6^, and during sexual development in the female *Anopheles* mosquito as oocysts (5,000-10,000 daughter sporozoites). Each of these population expansions serves to replenish parasite numbers following natural bottlenecks during transitions in the parasite life cycle. While the formation of daughter cells has been deeply analyzed during asexual blood stages^7^, the expansive process of sporozoite formation within the oocyst has remained relatively understudied^1^. Oocyst development begins when, after gamete fertilization and zygote formation in the blood bolus contained within the mosquito midgut lumen, a few ookinetes surviving crossing of the midgut epithelium encyst on the basal side of the epithelial layer just below the basal lamina^8-10^. Here, the oocyst grows extensively over the course of several days, undergoing prolific DNA and organelle replication, and around 7 to 8 days post-infection (dpi), following yet unknown cues, begins the highly coordinated process of daughter cell formation to replenish parasite numbers^11^. Pairing *de novo* cytoskeleton assembly with precise compartmentalization of essential organelles, this segmentation process culminates in the formation of thousands of infectious daughter sporozoites from a single oocyst (Fig. 1a). When mature, sporozoites are then released from the oocysts and travel to the mosquito salivary glands from where they are transmitted to humans when the *Anopheles* female next takes a blood meal. The sequence of events triggered by sporozoite injections causes the death of over half a million people (mostly young children in Africa) every year^12^ and is widely regarded as one of the strongest selective pressures to have shaped the human genome^13^.

**Fig. 1:**
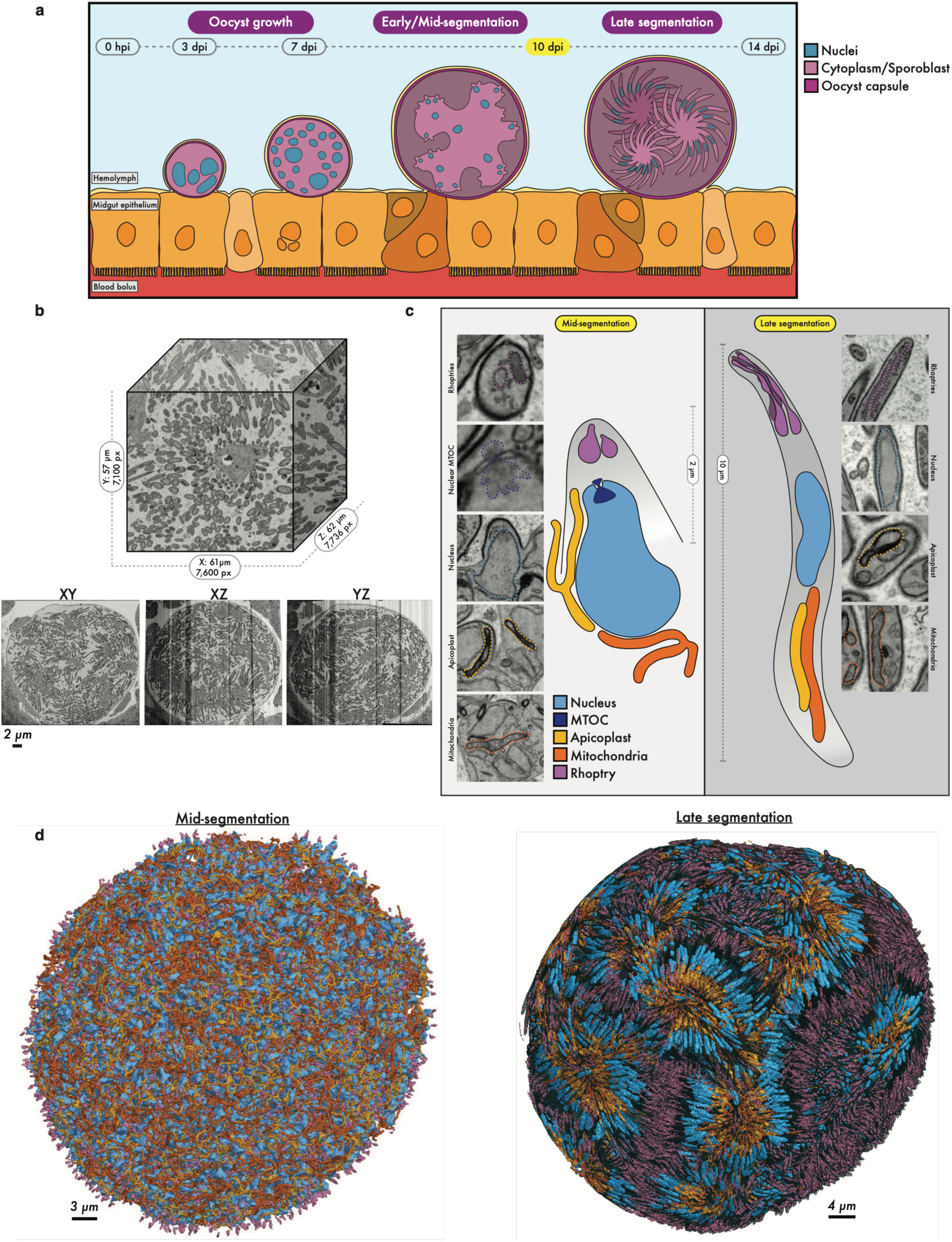
Enhanced FIB-SEM provides complete 3D reconstructions of *P. falciparum* oocysts during segmentation. **a,** Schematic of oocyst development in *An. gambiae* midguts, displaying overarching changes in parasite ultrastructure. Midguts infected with wild type *P. falciparum* NF54 parasites were collected from 20 mosquitoes and pooled at 10 dpi (days post infection). **b,** Schematic showing the full FIB-SEM volume for a single oocyst. Two volumes were acquired, capturing an oocyst at mid and late segmentation (with dimensions shown). Lateral (X-Y) and axial (X-Z and Y-Z) views of a small sub-region of the late segmentation dataset are shown, highlighting the isotropic resolution of the FIB-SEM volumes. px – pixel. **c,** Morphology of key organelles (dashed lines) as they appear in the raw FIB-SEM volumes during mid (left) and late segmentation (right), with schematic representations of daughter cell morphology. **d,** 3D reconstruction of segmented nuclei (*light blue*), nuclear microtubule organizing centers (MTOC) (*dark blue*), mitochondria (*orange*), apicoplasts (*yellow*) and rhoptries (*purple*) (inner membrane complex not shown) throughout the entire oocyst volume at mid (left) and late segmentation (right). Note that the nuclear MTOCs are no longer visible by late segmentation.

Despite their critical importance in disease transmission, to date, a detailed and complete ultrastructural overview of sporozoite formation in the oocyst has been lacking. Previous ultrastructural studies of *Plasmodium* replicative stages have focused primarily on blood stage schizogony, utilizing focused ion beam-scanning electron microscopy (FIB-SEM) to map inter-organelle interactions and structural changes at single digit nanometer resolution throughout the parasite volume, greatly expanding our knowledge of how 16-36 daughter merozoites are organized and formed at this stage^14-16^. Daughter cell formation in the oocyst is, however, several orders of magnitude more complex than blood stage schizogony and it remains unclear how conserved these processes are given the sheer difference in scale. Historically, studies by transmission electron microscopy of oocyst ultrastructure during segmentation have been largely limited to model species (e.g. the rodent parasite *Plasmodium berghei*) and information is constrained to what can be gleaned from single sections, preventing an understanding of how organelles interact through the entire cell volume^11,17-20^. Three-dimensional, whole cell ultrastructural studies have previously been precluded in the oocyst primarily owing to the significantly larger size of the imaged cell: a mature oocyst that has begun the process of segmentation is spherical and roughly 50 – 60 µm in diameter, equating to a total volume of about 113,000 µm^3^ (as compared to a schizont at ca. 100 µm^3^) which far exceeds the capabilities of most commercial FIB-SEM systems. Traditionally, imaging a cell of this size would necessitate a tradeoff between total acquisition volume preventing its complete reconstruction, or a dramatic decrease in resolution rendering the visualization of key ultrastructural features and inter-organelle interactions unreliable. Here, leveraging an enhanced FIB-SEM platform and a suite of custom-trained deep learning models, we present a complete reconstruction of the entire oocyst volume at nanometer resolution at two distinct time points in segmentation, capturing key ultrastructural changes essential for sporozoite formation and revealing new features of parasite biology at an unprecedented resolution.

### Reconstruction of sporozoite segmentation over time at nanoscale resolution

We infected *Anopheles gambiae* females with *Plasmodium falciparum* parasites, as this is the most important vector-parasite pairing in sub-Saharan Africa, and dissected infected midguts at different time points post-infection. As expected, we found significant heterogeneity in the timing of segmentation even within a single midgut, with oocysts showing different developmental stages. We therefore focused on 10 days post infection (dpi) as it represented a time point where several stages of segmentation were readily observable (Extended Data 1). We performed multiple rounds of x-ray microtomography at increasing resolution to identify two individual oocysts for FIB-SEM imaging (Methods). We chose an oocyst at mid-segmentation, where the nascent buds of the daughter cell cytoskeleton can be observed and organelle segregation is underway, and an oocyst very late in segmentation, where daughter sporozoites are almost fully elongated and organelle segregation is expected to be largely complete (Fig. 1a). We reasoned that comparison between these two time points would allow us to reconstruct the overall morphological changes oocysts undergo during segmentation and identify key organizational strategies utilized to ensure proper organelle segregation.

FIB-SEM was originally developed for research applications which did not necessitate large volume imaging and has only recently been adopted for the 3D imaging of biological specimens^21-24^. While allowing for isotropic single digit nanometer imaging, major barriers include slow imaging speed (capping the maximum possible image volume) and lack of long-term system stability which, due to the inherently destructive nature of FIB milling, can yield irreversible sample loss. Previous development of an enhanced FIB-SEM platform^2^, introducing hardware and software innovations, overcomes these challenges and opens the door for reconstructing the entire volume of *Plasmodium* oocysts at nanoscale resolution. The samples were imaged on a custom FIB-SEM system with a Zeiss Capella FIB column mounted at 90° on a Zeiss Gemini SEM^2^. Each block face was imaged by a 3.0 nA electron beam with 1.2 kV landing energy at 2 MHz. The X-Y pixel size was set at 8 nm. A focused Ga^+^ beam of 15 nA at 30 kV was strafed across the surface and ablated away 4 nm of the surface, and the newly exposed surface was then imaged again. Alternating rounds of ablation and imaging continued for about two weeks for each sample to capture the entirety of the oocysts. The raw imaged volume was subsequently post-processed with image registration and alignment followed by destreaking and smoothing (Methods).

With resultant image sizes of over 7,000 slices along each major axis (per dataset) and thousands of instances of features of interest in each slice (Fig. 1b), we reasoned that pure manual annotation of the two datasets would be infeasible. Instead, we leveraged machine learning workflows for image segmentation^3^. In brief, we successfully trained models *de novo* for six major organelle and structural features: nuclei, mitochondria, apicoplasts (parasite-specific metabolic organelles)^25^, microtubule organizing centers (MTOC), rhoptries (organelles used during host cell invasion)^26^ and inner membrane complexes (IMCs, a key structural component of the parasite cytoskeleton)^27^ in the mid-segmentation dataset (Fig. 1c, left). We further fine-tuned five of those models to accurately label the late-segmentation dataset, excluding the nuclear MTOCs which are no longer visible by this stage (Fig. 1c, right). Of note, *Plasmodium* lacks canonical centrioles, relying instead on the bipartite MTOC localized to the nuclear membrane^16,28,29^. Despite the abundance of pre-trained deep learning models to accurately label organelles (such as the mitochondria) in other eukaryotic cell types, the morphological divergence of *Plasmodium* in addition to the presence of parasite specific organelles (e.g., the apicoplast, IMC and rhoptries) necessitated a laborious training of custom models to accurately label our datasets (this process is described in detail in the methods section). In all, we were able to successfully reconstruct these structural features in both stages and counted 6,707 daughter cells in the mid-segmentation oocyst and 8,788 daughter cells in the late segmentation sample (Fig. 1d, Supplemental Video 1,2). The raw FIB-SEM data and resultant machine learning segmentations for both mid- and late segmentation are available through figshare (10.6084/m9.figshare.34918407 and 10.6084/m9.figshare.34925229, respectively)^30,31^.

### FIB-SEM imaging reveals distinct shifts in organelle morphology

These detailed reconstructions allowed us to observe key features of organelle morphology and their evolution between the two timepoints. At the onset of segmentation, the entire biomass of inherited organelles (nuclei, apicoplasts, and mitochondria) is contained within a central body bounded by the parasite plasma membrane, termed the sporoblast, from which all budding daughter cells will emerge (Fig. 1a). The daughter IMC, a core component of the parasite cytoskeleton that marks the boundaries of daughter sporozoites, is formed *de novo* at the onset of segmentation to provide the underlying structure of fully elongated, mature sporozoites^27^. In the mid-segmentation dataset, the cone-like structure of early IMC formation can be clearly observed, while by late segmentation this structure has elongated and undergone a significant increase in both volume and surface area (Fig. 2a-c) (Supplemental Video 3,4). The morphologies of many organelles shift dramatically between mid- and late segmentation as they are compartmentalized into the newly forming IMCs (Supplemental Video 5). Notably, the rhoptries, secretory organelles that are synthesized *de novo* and in sporozoites are essential for invasion of both mosquito salivary glands and human liver cells, undergo marked expansion and elongation over the course of sporozoite segmentation (Fig. 2d). In mid-segmentation, rhoptries are still immature and appear as apically positioned club-shaped organelles that closely resemble those in blood stage schizogony^15^. In daughter cells with more than one rhoptry, one organelle always appears less apically localized, less electron dense, and with a rounder shape (likely the equivalent of pre-rhoptries in blood stage schizogony), in line with previous observations by expansion microscopy in oocysts^15,32,33^ (Fig. 2d). By late segmentation, rhoptries are significantly elongated as well as increased in number, and micronemes (other organelles which are similarly involved in host cell attachment) can also be observed (Fig. 2d).

**Fig. 2:**
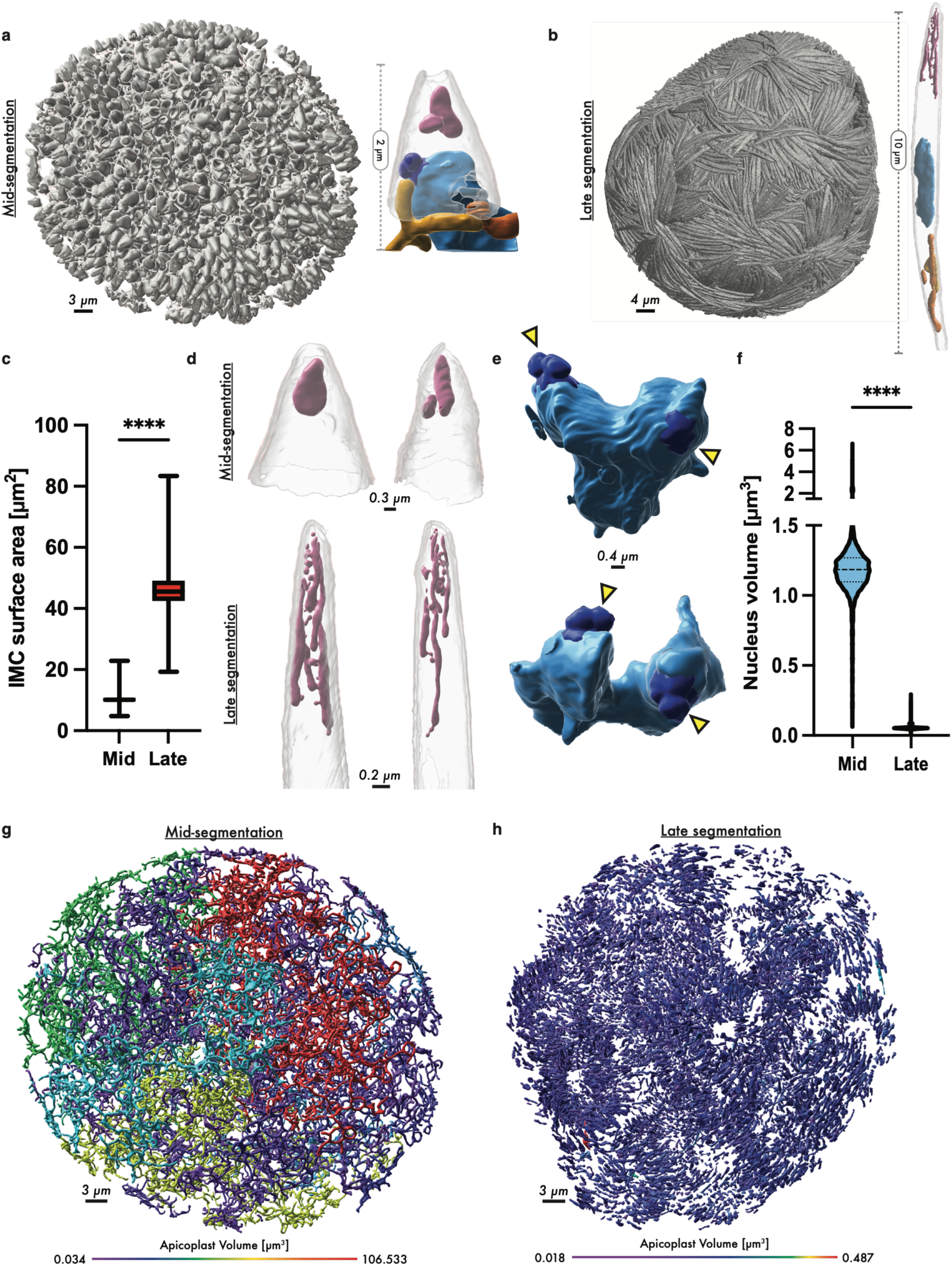
Organelle morphology shifts over the course of segmentation. **a,** 3D reconstruction of all 6,707 daughter inner membrane complexes (IMCs) during mid segmentation. Shown right is a crop of a single rendered IMC (*transparent*) containing all organelles: nuclei (*light blue*), nuclear MTOC (*dark blue*), mitochondria (*orange*), apicoplasts (*yellow*) and rhoptries (*purple*). **b, 3D** reconstruction of all 8,788 elongated IMCs during late segmentation. Shown right is a single rendered IMC. Colors as in a. **c,** IMCs elongate significantly in surface area as segmentation proceeds. Error bars denote 95% confidence interval. **d,** Rhoptry morphology shifts from typically 1–2 pear-shaped rhoptries per IMC in mid-segmentation (top) to multiple, elongated organelles in late segmentation (bottom). Colors as in a. **e,** Karyokinesis remains incomplete during mid-segmentation as evidenced by a small sub-population of nuclei that contain multiple nuclear MTOCs (*yellow arrowheads*). Colors as in a. **f,** Nucleus volume decreases significantly between mid and late segmentation and late segmentation nuclei are more homogenous in size. **g,** In mid-segmentation, apicoplasts have undergone intermediate scission and the 262 distinct plastid organelles vary widely in volume. **h,** Apicoplast scission is complete in late segmentation and the volumes of individual organelles occupy a tighter range. Differences in IMC surface area (µm^2^) and nuclei volume (µm^3^) between mid- and late segmentation were compared using Mann-Whitney two-tailed test (**c,f**). ****=P<0.0001.

In contrast, inherited organelles (nuclei, apicoplast, mitochondria) have already experienced large scale expansion by the time segmentation starts. We expected that karyokinesis would be complete by the start of cytokinesis, following multiple rounds of DNA replication and division earlier in oocyst development. However, in mid-segmentation, while cytokinesis is well underway, karyokinesis remains ongoing. Multi-lobulated nuclei can be readily observed, with the lobes extending into separate daughter IMCs (Fig. 2e). Karyokinesis is instead completed by late segmentation, where nuclei are largely uniform in size, occupying a region near the center of the elongating sporozoite (Fig. 2f, Extended Data 2a). As expected, the MTOC, which spans the nuclear membrane during mid-segmentation (Fig. 2e), is no longer discernible by late segmentation, when its role is complete.

In addition to the nuclear genome, *Plasmodium* parasites must also inherit at least one copy of the mitochondrial genome and a third distinct genome derived from the parasite-specific apicoplast. Both organelles are maternally inherited, and at the onset of oocyst development they appear as single, tubular structures, and undergo prolific expansion in organelle biomass prior to the onset of segmentation^34,35^. In the mid-segmentation oocyst, we capture this organellar expansion in its entirety, where both the mitochondrion and apicoplast are observed as convoluted, highly branched structures that stretch throughout the oocyst volume (shown in Fig. 2g for the apicoplast, and in Extended Data 3a,b for the mitochondrion). Interestingly, we found that the apicoplast at this time point is not a single, contiguous organelle but is rather comprised of 262 discrete organelles of varying sizes (Fig. 2g, Extended Data 3c). This suggests that this organelle has completed its expansion at this stage and is starting to undergo scission. In late segmentation, the apicoplast has completed scission and can be observed as a tubular organelle in close proximity to the mitochondrion (Fig. 2h). The mitochondrion, which at this time point is observed as a more elongated, tubular organelle, has instead yet to undergo the final scission event and remains attached to the sporoblast, as also confirmed by expansion microscopy (Supplemental Video 6, Extended Data 4a-c). Of note, we could not count the number of individual mitochondria during mid-segmentation due to technical difficulties with labeling the large branching structure in a contiguous manner, with the dataset providing an overestimate of mitochondria numbers due to single surfaces mis-segmented as separate objects (see Methods).

### Segmentation at scale is remarkably accurate and synchronous

To assess the fidelity and synchronicity of segmentation, we examined the organellar contents of all daughter IMCs across both cells. To this end, we used the IMC boundaries to computationally define the bounding edges of each daughter cell and, for the mid-segmentation dataset, superimposed a convex hull onto the IMC to approximate a closed shape (Methods, Extended Data 5a,b). We found that at mid-segmentation roughly 99% of daughter IMCs contain one or more rhoptries, a nucleus, and corresponding nuclear MTOC. Moreover, 88% of IMCs contain at least part of an apicoplast branch, while the mitochondrion lags behind (only 4% of IMCs) (Fig. 3a). This is in line with previously described dynamics in the blood stages of parasite development, where it was found that compartmentalization and scission of the apicoplast precedes mitochondrial segmentation^15,16^. By late segmentation, 99% of daughter cells contain rhoptries, apicoplasts, and mitochondria, while 94% contain a nucleus (Fig. 3b). These results indicate a high degree of fidelity in segmentation, where most daughter cells receive a complete set of essential organelles, a particularly striking finding given the thousands of daughter cells that are simultaneously formed in a single oocyst. Such degree of concordance in the organellar contents of each IMC indicates that segmentation proceeds in a highly synchronous manner, supported by the tight, homogenous distribution of IMC lengths at both mid- and late segmentation (Fig. 2c). Importantly, and contrary to previous assumptions, we conclusively visualize and mathematically confirm (see Methods) a single, interconnected sporoblast at both mid- and late segmentation, possibly facilitating the signaling events underlying the coordinated timing of daughter cell formation (Fig. 3c).

**Fig. 3:**
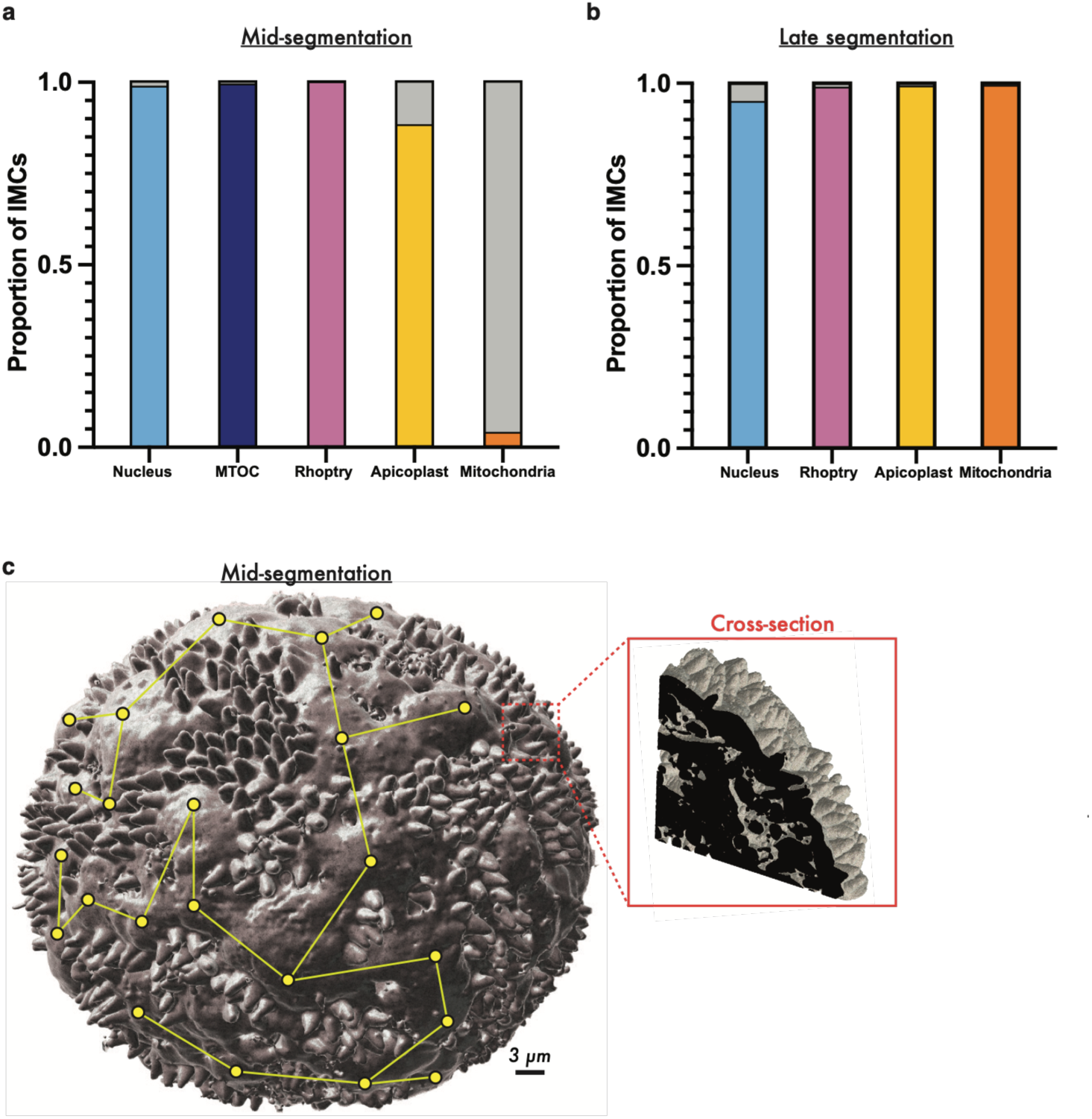
Organelle segregation is accurate and synchronous. **a–b,** The majority of IMCs contain all organelles at mid-segmentation except the mitochondria (**a**), whose segregation lags behind, but is complete by late segmentation (**b**). **c,** The sporoblast (connected by green lines) is one contiguous structure in mid-segmentation. Inset shows a region in cross-section.

### Key inter-organelle interactions may ensure fidelity of segregation

In addition to changes in organelle morphology over the course of segmentation, we also noted several inter-organelle interactions. In mid-segmentation, we observed a distinct association between the distal tips of apicoplast branches and the extra-nuclear portion of the MTOC, with the two organelles appearing to be in direct contact (Fig. 4a,b). While this association has been previously observed in *Toxoplasma gondii^36^*, *Sarcocystis neurona^37^*, and the blood stages of *Plasmodium* development^16^, here we could test its statistical significance given the large number of daughter cells formed in a single oocyst (6,707 in this dataset). By measuring the actual distance from each MTOC in the dataset to the nearest apicoplast and comparing it to the distance of 1,000 iterations of mock completely spatially randomized (CSR) distributions of MTOCs to the nearest apicoplast in each iteration (Methods), we demonstrated a non-random clustering of the two structures (Fig. 4c). No association was instead found between the MTOC and the mitochondria, despite them also forming a large branching structure that stretches throughout the oocyst volume (Fig. 4c). In late segmentation instead the mitochondria are closely associated with the apicoplasts (Fig. 4d), with 98% of apicoplasts 0.0 µm away from the nearest mitochondrion (Fig. 4e). This direct association likely facilitates the exchange of key metabolic components whose synthesis is shared between the two organelles^38^.

**Fig. 4:**
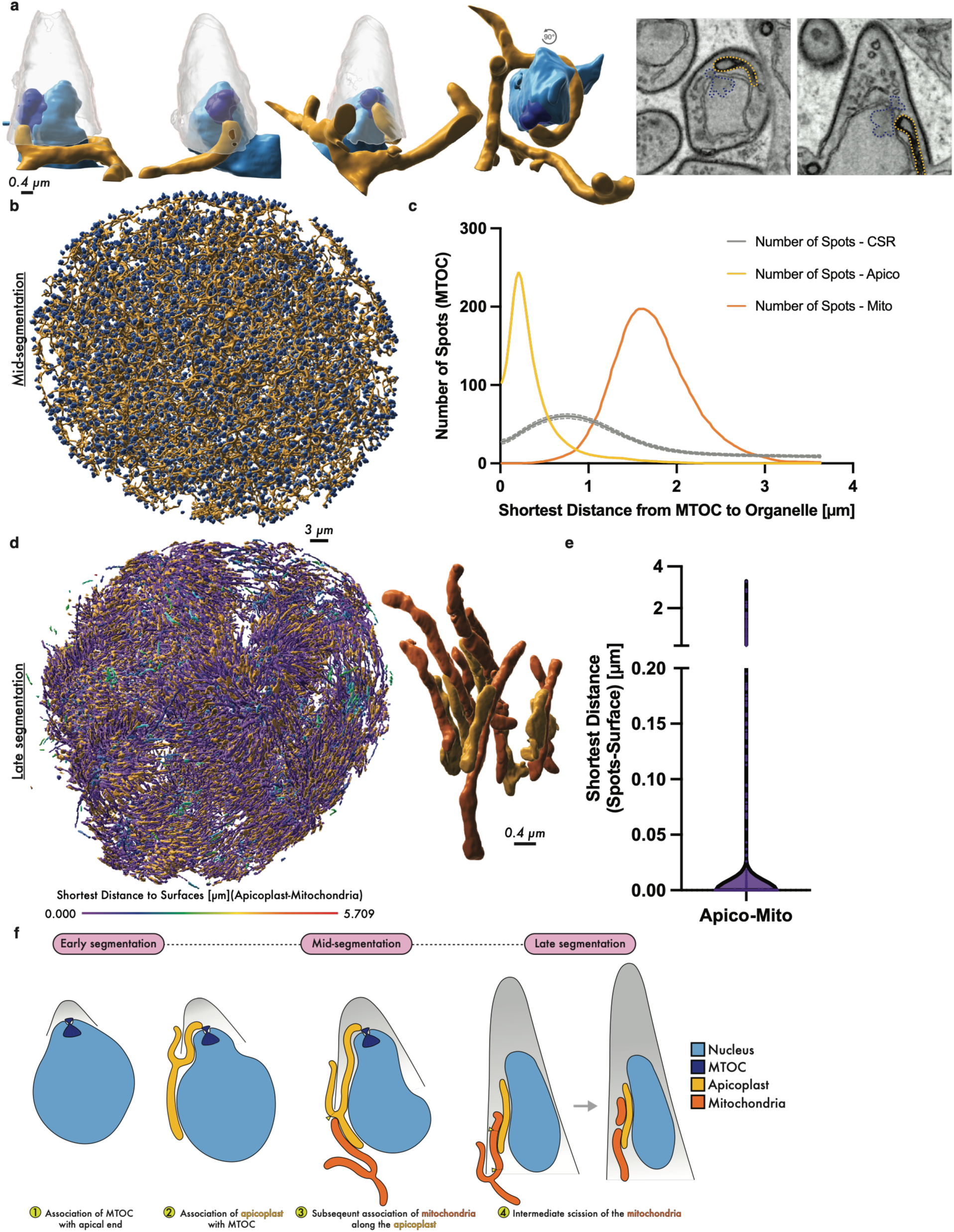
Inter-organelle interactions may facilitate accuracy during segregation. **a-b,** Apicoplasts (*yellow*) associate tightly with the nuclear MTOCs (*dark blue*) during mid-segmentation. (**a)** shows a 3D reconstruction and raw EM of a zoomed in subset of individual instances of this interaction (IMC, *transparent*; nuclei, *blue*), while (**b**) shows the entire oocyst (same colors as **a**). **c,** Tight association between apicoplasts and MTOCs, as measured by the shortest distance between each MTOC and the nearest apicoplast (*yellow line*) than would be expected by random chance in 1,000 spatially randomized distributions of MTOCs (*grey line, upper and lower limits dashed lines*). No association is instead found between the mitochondria and the MTOCs (*orange line*). **d,** Mitochondria and apicoplasts are in direct contact during late segmentation. 3D reconstruction of all apicoplasts (*yellow*) and mitochondria (*color-coded by shortest distance to nearest apicoplast*) and zoomed in subset of apicoplasts (*yellow*) and mitochondria (*orange*). **e,** Median distance between apicoplasts and mitochondria is 0.0 µm (0.0 µm 25^th^ quartile; 0.0 µm 75^th^ quartile). **f,** Stepwise model of inter-organelle associations during segmentation.

Our data however revealed a distinct difference in the fidelity of apicoplast and mitochondrial segmentation. Late in sporozoite formation, when 99% of daughter cells contain at least one copy of each organelle, we found more than one mitochondrion in 36% of cells, 96% of which possessed three or more mitochondria. By contrast, just 1.4% of daughter cells contained more than one apicoplast, and a mere 0.057% had three or more (Extended Data 6). This may be a direct consequence of the differing methods of inter-organellar associations: a singular point of association between the apicoplast and MTOC compared to the multiple points of association between the mitochondria along the length of the apicoplast, that might affect the precision of cellular counting.

Based on these observations, our hypothesis is that tight inter-organelle associations at different stages of segmentation may represent a cohesive organizational strategy to ensure each daughter cell receives at least one apicoplast and mitochondrion per nucleus. In this model, the earlier MTOC-apicoplast association ensures correct segmentation of plastids, with the mitochondrion hitching a ride later via its close association with the apicoplast (Fig. 4f).

### Nanoscale resolution reconstructions unveil novel sporozoite biology

In the late segmentation oocyst, the fine nanometer resolution of our imaging dataset enabled us to visualize several ultrastructural features not previously described in the literature. In a significant proportion of sporozoites, we observed the presence of a distinct membrane-bound protrusion emanating from a clear gap in the parasite cytoskeleton (Fig. 5a,b, Supplemental Video 7). These compartments contain membranous whorls and cytoplasm and, in some instances, they enclose fragments of organelles such as the mitochondria. From quantification of a subset of parasites we estimate that 45% of sporozoites excrete these blebs, the vast majority of which (91%) are observed along the convex curve (Fig. 5c). These structures do not resemble standard artifacts of electron microscopy sample preparation, as they are surrounded in all cases by a continuous and undisrupted plasma membrane which is not typically observed in instances of aberrations due to sample handling (Fig. 5a). Instead, the corresponding gap in the parasite cytoskeleton paired with the uninterrupted plasma membrane suggests an active remodeling process that modifies the parasite cytoskeletal structures to facilitate extrusion of these membranous compartments (Fig. 5d).

**Fig. 5:**
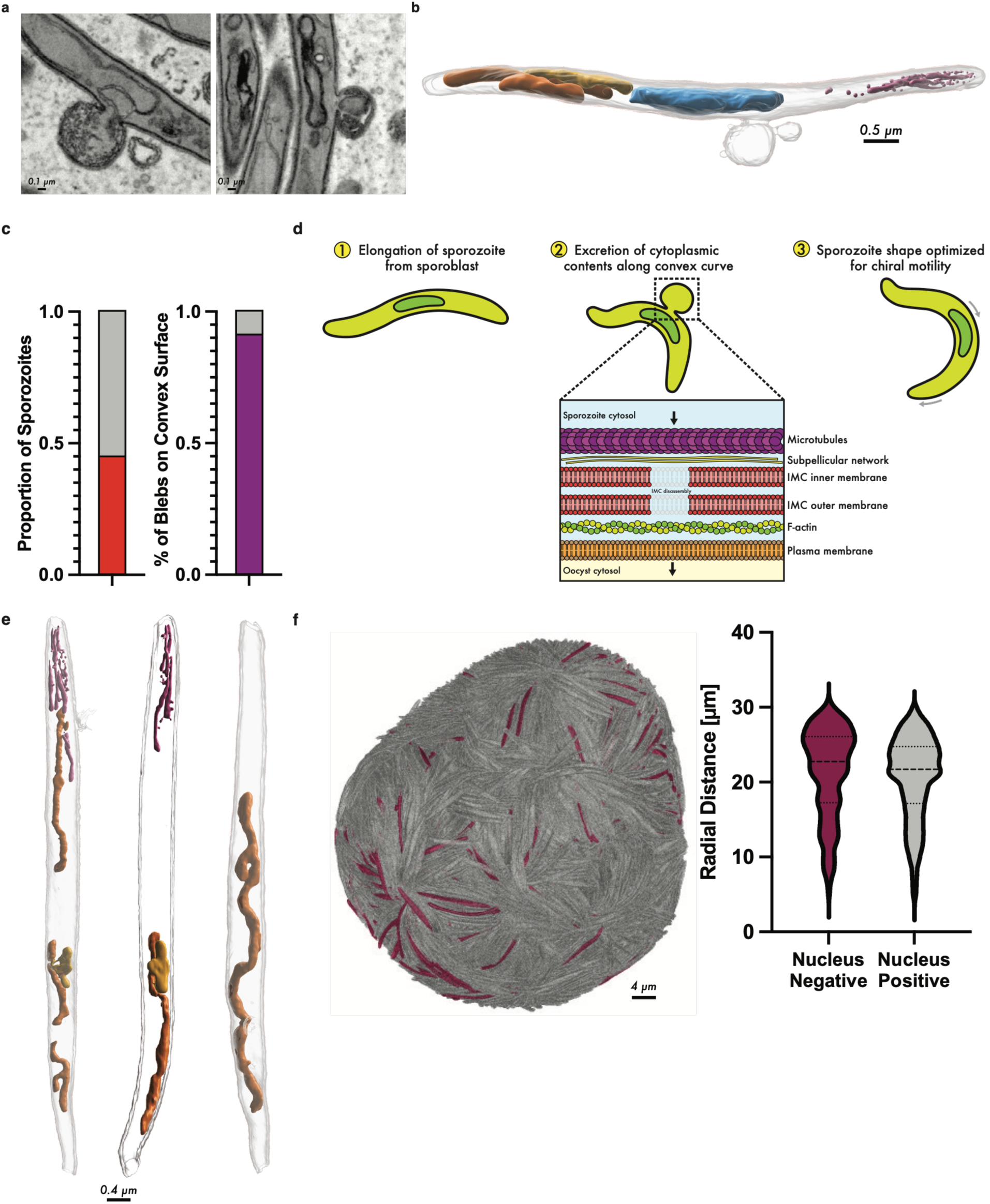
FIB-SEM reveals novel sporozoite biology. **a-b,** Sporozoites excrete membrane bound-cytoplasmic compartments from discrete breaks in the IMC. (**a**) Raw EM highlighting the distinct break in the electron-dense IMC from which these membrane-bound structures are extruded, showing they contain cytoplasm and membranes (*left*) and occasionally fragments of organelles such as the mitochondria (*right*). (**b**) 3D reconstruction (plasma membrane, *transparent*; apicoplast, *yellow*; mitochondria, *orange*; nuclei, *blue*; and rhoptries, *purple*). **c,** From quantification of a subset of 100 developing sporozoites, 45% could be observed with these membranous protrusions and of those, blebs were observed along the convex curve of the sporozoite 91% of the time. **d,** Bleb formation model. **e,** 3D reconstruction of a representative subset of sporozoites in late segmentation that completely lack a nucleus despite possessing other key organelles. Same colors as **a**. **f,** Anucleate sporozoites are observed through the oocyst volume, 3D reconstruction (nucleated sporozoites, *white*; anucleate sporozoites, *burgundy*). Quantification of distribution of nucleated and anucleate sporozoites as measured by radial distance from the center of the oocyst.

As hinted at earlier, we also observed the presence of a sub-population of sporozoites that completely lacked a nucleus, despite the advanced nature of segmentation and the presence of spare nuclei in the sporoblast (Fig. 5e, Extended Data 7a). These anucleate ‘zombie’ sporozoites often contain all other organelles (apicoplast, mitochondrion and rhoptries), although we also observed several instances of daughter sporozoites that only contained mitochondria and were devoid of all other organelles. Anucleate sporozoites account for about 5% of the total number (446 out of 8,788) and are distributed randomly throughout the oocyst volume (Fig. 5f). We hypothesize that this segregation error in sporogony is the consequence of an innate strategy wherein the oocyst forms daughter cells as rapidly as possible, but at the cost of a small degree of waste where excess amounts of nuclei and other organelles are formed but are never correctly segregated into daughter cells. Supporting this hypothesis, we also observed remnants of all other organelles within the residual sporoblast structure (Extended Data 7a-c).

## Discussion

By focusing on a key replicative stage of *P. falciparum* development that has long escaped detailed ultrastructural study, our work provides a comprehensive reconstruction of subcellular features involved in the formation of sporozoites, the parasite stage infectious to humans. Through this work, we examine two distinct time points in segmentation and observe a large conservation in the timing of organelle segregation and key inter-organelle interactions with blood stage parasites, despite the vast difference in scale of these segmentation processes (tens of merozoites versus thousands of sporozoites formed from a single round of division). We demonstrate that karyokinesis is largely complete relatively early during the segmentation process, although surprisingly, a small subpopulation of nuclei appears to still be in the process of dividing as demonstrated by their greater volume (≥3 µm^3^ on average) and the presence of two MTOCs in a single nuclear body (Fig. 2e). We reveal that the parasite-specific metabolic organelle, the apicoplast (and likely the mitochondrion, although we cannot conclusively prove this) has already undergone the early stages of scission at mid-segmentation, while at this stage the rhoptries have started forming but are not yet mature. By late segmentation, the apicoplast has fully divided, the rhoptries have fully matured, but the mitochondria remain clearly tethered to the sporoblast, proving this is the last organelle to be segregated into sporozoites (Extended Data 4). These observations support previous findings from single cell RNAseq showing that expansion of the apicoplast and mitochondrion occurs first during oocyst development, followed by rhoptry biogenesis at the start of segmentation^39^.

The observed organelle dynamics reveal a stepwise model likely aimed at ensuring the fidelity of organelle segmentation, with parallels to observations in asexual blood stages^15,16,32,40^. Nuclear division occurs first, with the MTOC-containing portion of the nucleus oriented towards the apical end of the elongating IMC, likely through a second interaction with the newly forming apical tip of the parasite. Subsequently, the apicoplast associates with the MTOC, followed by the mitochondrion associating along the length of the apicoplast. We hypothesize that these sequential associations represent an overarching organizational strategy to ensure each daughter cell not only receives a single copy of each organelle but also at least one copy of the three distinct genomes (nuclear, apicoplastid, and mitochondrial). Given the conservation of these associations at the larger scale of the oocyst, it is tempting to speculate that similar strategies may be utilized during daughter cell formation in liver stage schizogony where thousands of daughter cells are also formed.

Organelle partitioning is a complex, multi-step process. Previous models of this process describe two overlapping phases with a host of mechanistic and molecular insights: 1) organelle movement mediated by the canonical actomyosin complex^41-43^, and 2) organelle retention governed by additional contacts with molecular chaperones and adapters^44^. Based on our work and integrating previous observations^16,36,37,45^, we propose a third phase: organelle counting mediated by interactions with the MTOC which acts as a proofreading mechanism to ensure the fidelity of segmentation. Proofreading may have an outsized impact in oocysts which must organize the organellar contents of orders of magnitude more daughter cells than *T. gondii* (2 tachyzoites) or *P. falciparum* blood stages (16-36 merozoites). Indeed, recently in *T. gondii*, the apicoplast-MTOC association was identified as being mediated at least in part by apicoplast partitioning proteins (AAP1/3), depletion of which resulted in apicoplast segregation defects^46^. Of note, the molecular components governing organelle segregation in the oocyst remain largely unknown. While it is likely some proteins identified in the blood stages of development perform conserved functions, we also hypothesize that oocysts utilize stage-specific proteins to help account for the differences in division dynamics. Specifically, components essential for organelle segregation in blood stage schizogony lack evidence of expression during the oocyst stages of development^39,47^. As highlighted above, combining high-resolution microscopy with transcriptional data at the single cell level thus offers the opportunity to gain novel insight into the specific processes mediating organelle biogenesis and fission in nascent sporozoites.

Among unexpected findings, we visualized membrane-bound compartments that protrude from discrete openings in the parasite cytoskeletal structures. The content of these blebs is largely cytoplasmic but occasionally includes fragments of organelles (e.g. mitochondria). We posit that these structures represent a means for the cell to excrete waste and refine cell shape during sporozoite formation in preparation for downstream motility, which is critical for sporozoite egress from the oocyst and subsequent infectivity to a human host (Fig. 5d). Indeed, the asymmetric localization of these blebs almost exclusively along the convex curve of the IMC supports the theory that their excretion may play a role in maturation of motile daughter cells, as sporozoites are known to exhibit chiral, helical motion when gliding through three-dimensional tissue environments^48,49^. Additional studies are needed to define the molecular factors governing the cytoskeletal remodeling and formation of these structures, and their role in sporozoite maturation.

Our data reveal that the oocyst exhibits a high degree of accuracy in the process of forming thousands of daughter cells, although occasional mistakes occur. The identification of a small subpopulation of fully elongated sporozoites that lack a nucleus despite the presence of numerous residual nuclei in the sporoblast, indicates that the oocyst does not possess an innate mechanism to ensure a perfect one-to-one pairing of daughter cells formed to copies of organelles produced, nor an abortive checkpoint for IMC elongation. Instead, commitment to segmentation represents an irreversible cell fate likely initiated once the oocyst has produced a significant biomass of organelles. Despite these errors, the overall rate of accuracy during segmentation is strikingly high, with 95% (8,342/8,788) of IMCs having all essential organelles. Given the high number of correctly segmented sporozoites, parasites can tolerate this small degree of error without compromising their transmissibility. Thus, parasites play a balancing act during segmentation, where rapid expansion and organelle replication are prioritized during the growth phase yielding an excess of organelles, followed by tightly coordinated organizational strategies to ensure that daughter cell formation is as accurate as possible. Future studies are needed to better understand the limiting factors that constrain the number of organelles and daughter cells that are ultimately formed within each oocyst, and whether this complex, yet beautiful, process can be targeted for new transmission blocking strategies.

## Supporting information

Supplemental Video 1: 3D reconstruction of all organelles in mid-segmentation

Supplemental Video 2: 3D reconstruction of all organelles during late segmentation

Supplemental Video 3: Mid-segmentation FIB-SEM data overview

Supplemental Video 4: Late segmentation FIB-SEM data overview

Supplemental Video 5: Shifting IMC and organelle morphology during segmentation

Supplemental Video 6: Mitochondria remained tethered to the sporoblast in late segmentation

Supplemental Video 7: Maturing sporozoites excrete membrane-bound compartments

Supplemental Information Guide

## Methods

### Mosquito rearing

*Anopheles gambiae* (G3 strain) mosquitoes were housed in cages in an insectary maintained at 27°C, 70-80% relative humidity, and a 12-hour light-dark cycle. Mosquitoes were provided with water and 10% w/v glucose (Sigma-Aldrich; St. Louis, MO) solution *ad libitum*, and fed on human blood (Research Blood Components; Boston, MA) once a week for colony maintenance. Mosquitoes used in experiments were taken from the colony as pupae and allowed to eclose in a cage; adults were provided with water and 10% w/v glucose solution *ad libitum* until time of experimentation.

### *Plasmodium falciparum* culturing and gametocyte induction

*Plasmodium falciparum* NF54 was used for all experiments. Asexual stage parasites were maintained in RPMI 1640 with L-glutamine and 25 mM HEPES buffer (Corning; Corning, New York, USA) supplemented with 10 mg/L hypoxanthine, 0.2% sodium bicarbonate, and 10% heat-inactivated human serum (Interstate Blood Bank; Memphis, TN) and 5% hematocrit with human red blood cells (Research Blood Components; Boston, MA). Parasite cultures were kept in incubators at 37°C and gas composition of 5% O_2_, 5% CO_2_, and balanced N_2_ for up to two months in line with established protocols^50^. Gametocytogenesis was induced by increasing parasitemia to >2% and incubating for 14–20 days with daily media changes to accumulate mature stage V gametocytes, following established protocols^51^.

### *Anopheles gambiae* infection with *Plasmodium falciparum*

4–7-day old *An. gambiae* females were fed with stage V *P. falciparum* gametocyte cultures maintained at 37°C in custom glass, water-jacketed membrane feeders. Mosquitoes were allowed to feed for up to 60 minutes before being placed in a custom-made glovebox for biosafety containment (Inert Technology; Amesbury, MA). Any mosquitoes that did not appear fully engorged were removed. Blood fed mosquitoes were provided water and 10% glucose solution *ad libitum*.

### Infected midgut dissections

At 10 days post-infection, mosquitoes were aspirated into 80% ethanol, incubated at - 20°C for 10 minutes and then transferred to 1X PBS at 20–22°C for dissection. Midguts were dissected into 1X PBS and immediately transferred into a fixative solution of 2.5% paraformaldehyde, 5% glutaraldehyde, 0.06% picric acid in 0.2M cacodylate buffer and spun down briefly to fully submerge tissues in fixative. Fixed tissues were submitted to the Harvard Medical School Electron Microscopy Core for staining and initial embedding.

### Electron microscopy sample preparation

Samples were prepared using a Reduced Osmium-Thiocarbohydrazide-Osmium (rOTO) staining protocol. Fixed samples were washed in 0.1M cacodylate buffer, twice in water, and then postfixed in 1% osmium tetroxide/1.5% potassium ferrocyanide in water for 1 hour followed by 1% thiocarbohydrazide in water, and 1% osmium tetroxide in water. Samples were then washed twice in water followed by a single wash in 50 mM maleate buffer (pH 5.15). Next, the samples were incubated overnight in 1% uranyl acetate in maleate buffer, followed by one wash in maleate buffer, and two washes in water. Samples were then dehydrated via an ethanol gradient (50%, 70%, 90%, 100%, 100% ethanol) for 10 minutes at each concentration. After dehydration, samples were placed in propylene oxide for one hour and then infiltrated in a 1:1 mixture of propylene oxide and PELCO Eponate 12 overnight. Samples were then embedded in PELCO Eponate 12 with 31% Nadic methyl anhydride, 17.4% dodecenyl succinic anhydride, and 3% benzyldimethylamine and polymerized at 60°C for 48 hours. For quality control, ultrathin sections (80 nm) were cut on a Reichert Ultracut-S microtome, and sections were picked up onto copper grids and imaged in a JEOL 1200EX transmission electron microscope equipped with an AMT 2K CCD camera.

### FIB-SEM imaging

Prior to FIB milling, samples were re-mounted on a copper stud and subjected to serial rounds of ultramicrotome trimming (Leica EM UC7) and X-ray tomography (Zeiss Xradia 510) to locate oocysts representing mid- and late segmentation within the resin block by oocyst size and surface smoothness. A final “tab” of dimensions ca. 100 µm x 100 µm x 65 µm was trimmed from the block containing the region of interest. The trimmed block was sputter coated with 10 nm gold and 100 nm carbon film thickness (Gatan 682 PECS). Samples were loaded onto a customized FIB-SEM, consisting of a Zeiss Gemini 450 Field Emission SEM and a Zeiss Capella FIB column oriented at 90 degrees to the SEM beam. Details of the enhanced FIB-SEM system were previously described^2^. Each block face was imaged using a scanned electron beam with 1.2 kV landing energy and 3.0 nA electron dose. Images were acquired at 2 MHz readout rate (0.5 µs pixel dwell time), and 8 nm pixel size. The sample surface was ablated using a 15 nA Ga+ source at 30kV acceleration voltage. Actual ablation time and gallium current was adjusted to maintain an average 4 nm milling rate per image. Successive images were binned 2x in Z, producing isotropic 8 nm pixel size datasets. Alternating rounds of ablation and imaging continued for about two weeks to complete FIB-SEM imaging of one sample. Resultant SEM image slices were registered with respect to each other using a python implementation of the SIFT algorithm with RANSAC solver and a regularized affine transformation model (available at https://github.com/gleb-shtengel/FIB-SEM). After alignment, FIB-SEM images were subject to Fourier-based filtering (MATLAB R2024b) to minimize streaking artifacts due to uneven sample hardness during Ga^+^ beam milling^52^.

### Machine learning-based segmentation of FIB-SEM data

Segmentations of inner membrane complex (IMC), nuclei, mitochondria, apicoplasts, rhoptries, microtubule organizing centers (MTOC), and capsules were produced using custom machine learning models generated with Empanada (Empanada Version 1.2) implemented in Napari^3^. An average of 30 to 150 image patches (512 x 512 pixels each) were manually annotated to generate ground truth datasets for each round of training per organelle class (on average 2-7 rounds of training), and all models were trained for an average of 7,000 iterations in each round of training. Model performance was considered sufficient with an Intersection over Union (IoU) score of 0.7 or greater and was assessed manually by visual congruence between predicted labels overlaid on the raw EM data. When necessitated by RAM limitations during model inference generation, data subsets were generated from the full MRC raw data file using the ‘newstack’ command in IMOD (IMOD Version 5.1)^53^. Segmentations generated on these subsets were exported from Empanada as .zarr files and later stitched back together using custom Python scripts.

Single-class segmentation models were trained for nuclei, apicoplast, mitochondria, rhoptries, and capsule using the Panoptic DeepLab architecture. Distinct morphological changes of organelles between mid- and late segmentation timepoints necessitated additional rounds of fine-tuning and manual ground truth annotation to adapt each mid-segmentation model to recognize and accurately label the late segmentation dataset. As such, two trained models (a mid- and a late segmentation model) were generated for each of these organelle classes. Models were applied to the entire dataset (subset into halves to facilitate memory usage) using default inference settings in Empanada.

Separate instance segmentation models were trained for the IMC and MTOC. One trained model was generated for the MTOC as these structures were absent by late segmentation; two trained models were generated for the IMC. The MTOC and mid-segmentation IMC models were applied to the entire dataset (subset into halves) using default inference settings in Empanada. The late segmentation IMC model was applied to the entire dataset (subset into sixteenths) using ortho-plane settings, where the inference is performed independently on the three principal planes (XY, YZ, and XZ) before a final segmentation is produced based on a two of three consensus.

Cytoplasmic droplets were manually labeled in a subset of daughter cells in the late segmentation dataset using Napari and required ca. 30 hours for each daughter cell.

Sporoblast segmentation was performed using MATLAB (MATLAB R2024b). An intensity threshold was empirically determined to isolate sporoblast from background (35100 and 34950 for the mid- and late segmentation datasets, respectively), followed by a capsule mask (produced by performing image closing and binary fill holes operations on the capsule segmentations) to remove features outside of the oocyst. Image opening was used to remove spurious small objects in the sporoblast segmentation.

### Segmentation proofreading

3D surfaces were rendered for each segmentation type in Imaris (Imaris 10.2.0; Oxford Instruments) with absolute intensity thresholding, surface smoothing (0.016 µm), and the application of organelle-specific size filters to help exclude small mis-segmentation artifacts. Rendered surfaces were then overlaid over the raw FIB-SEM data to allow for proofreading of segmentation results. Proofreading of labels included application of additional volumetric filters to eliminate very small segmentation artifacts in addition to manual correction, including deletion of larger segmentation artifacts, merging of overlapping separate surfaces, and splitting of erroneously merged surfaces for every surface in a given organelle class. For the IMC in late segmentation dataset, custom MATLAB code was used instead of Imaris to remove over-segmentations.

Apicoplast continuity and surface merging was performed manually in the mid-segmentation dataset to mitigate mis-segmentation artifacts. Similar proofreading for the mitochondria in mid-segmentation could not be performed due to the infeasibility of manually proofreading the >6,000 mitochondrial surfaces.

### Statistics of subcellular features

Individual and cumulative organelle statistics (e.g. volume) were derived in Imaris. To calculate the proportion of daughter cells in each dataset that contained a specific organelle type, custom MATLAB code was used to produce cutouts of each organelle class within a bounding box of each unique IMC label. Daughter cells were deemed to contain an organelle if any volume of that organelle was found within a 3D hull based on the IMC segmentation at each stage: at mid-segmentation, “within” the IMC was defined as overlapping with a mask produced from the convex hull of each IMC label; at late segmentation, a mask created using image closing and binary fill holes was used. This overlap analysis was done using custom codes in MATLAB (mid-segmentation) and Python (late segmentation).

Late segmentation daughter cells initially found to have no nuclei were manually validated as such using a custom MATLAB script to facilitate visualizing raw EM data overlaid with IMC labels of interest.

Each MTOC in the dataset was represented as a simplified 3D shape (known in Imaris as Spots), by finding the center of mass (COM) in the three principal axes and these COM coordinates were used to generate spherical representations of all MTOCs. Using these MTOC Spots, the shortest distance from each Spot to the nearest 3D apicoplast surface was measured throughout the entire dataset (Imaris). To assess whether this association was statistically meaningful or likely due to random crowding effects, Imaris Vantage was used to generate 1,000 simulations of completely spatially randomized (CSR) distributions of MTOC Spots throughout the volume of the oocyst excluding area within the apicoplast surface. For each simulation, the shortest distance was calculated from each CSR MTOC Spot to the nearest 3D apicoplast surface and subsequently plotted against the actual MTOC to apicoplast distances. Additionally, to assess whether this observation was specific to the apicoplast, the shortest distance was calculated from each MTOC Spot to the nearest mitochondrial surface. Statistical analyses were performed in GraphPad Prism 11.0 (GraphPad Software Inc.; La Jolla, CA) with a Mann-Whitney test and by calculating effect size *r*.

Sporozoite blebs were quantified manually in a randomly selected subset of 100 IMC cutouts in the late segmentation dataset.

Sporoblast segmentation was done using MATLAB. Each z-slice of the EM volume was Gaussian filtered (sigma = 3) before an empirically determined intensity threshold was applied to isolate sporoblast from background (35100 and 35180 for the late and mid segmentation datasets, respectively), then a capsule mask (produced by performing image closing and binary fill holes operations on the capsule segmentations) was applied to remove features outside of the oocyst. Custom python code was used to assess sporoblast connectivity by slab-wise running 3D connected-component analysis with 26-connectivity on the sporoblast mask. Manual proofreading of the retained largest connected component validated that the sporoblast forms one continuous volume, and any smaller components that were identified were determined to be spurious over-segmentations from the initial intensity threshold.

### Data availability

The raw FIB-SEM data and machine learning segmentations that support the findings of this study are available in figshare with the identifier(s) [https://doi.org/10.6084/m9.figshare.34918407 and https://doi.org/10.6084/m9.figshare.34925229]^30,31^. Data are viewable through a Neuroglancer link provided on each figshare page allowing for visualization of the data without specialized computational requirements.

### Code availability

The custom code used in this study will be available upon publication on Zenodo. Access prior to publication for replication or verification can be requested from Flaminia Catteruccia.

## Acknowledgements

FIB-SEM imaging and data analysis was performed in collaboration with the Advanced Imaging Center (AIC) at Janelia Research Campus, a facility generously supported by the Howard Hughes Medical Institute (HHMI). We thank Maria Ericsson and Anja Nordstrom of the Harvard Medical School Electron Microscopy Core Facility for EM sample preparation; Stephan Kraemer at the Harvard Center for Nanoscale Systems for preliminary imaging; and Malhar Khushu for helpful discussions. F.C. is funded by the Howard Hughes Medical Institute (HHMI) as an Investigator and by the National Institutes of Health (NIH) grants R01AI148646 and R01AI153404.

## Contributions

E.W.D. and F.C. conceived the study. E.W.D. performed sample collection. J.A. performed FIB-SEM imaging. E.W.D., E.H., and J.A. performed model training. E.W.D., E.H., J.A., W.R.S., and J.D.D. performed data analysis and visualization. T.-L.C. and F.C. provided supervision and oversaw experiments. E.W.D. and F.C. wrote the original manuscript draft. E.W.D., E.H., J.A., W.R.S., J.D.D., T.-L.C., and F.C. reviewed and edited the manuscript. All authors approved the final manuscript.

## Extended Data

**Extended Data 1:**
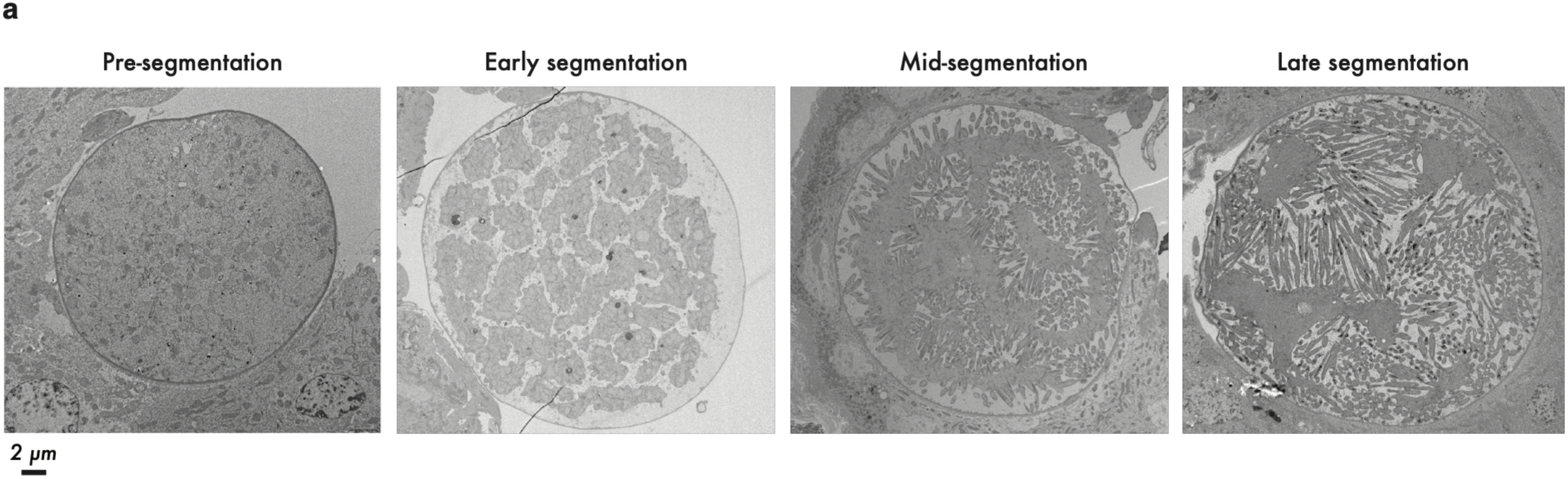
*P. falciparum* oocysts exhibit significant heterogeneity in development. **a,** Oocysts at 10 days post-infection can be observed at a diversity of developmental timepoints ranging from pre-segmentation (prior to formation of daughter cytoskeletal structures) to late segmentation (where sporozoite elongation is almost complete). Ultrastructural observation of oocyst stage was based on the presence/absence of the inner membrane complex (IMC) which is formed *de novo* during segmentation, and the subsequent length of the IMC and whether organelles could be observed primarily within the sporoblast or within the boundaries of the IMC. We selected an oocyst at an intermediate stage between early and mid-segmentation and one at very late segmentation for FIB-SEM imaging.

**Extended Data 2:**
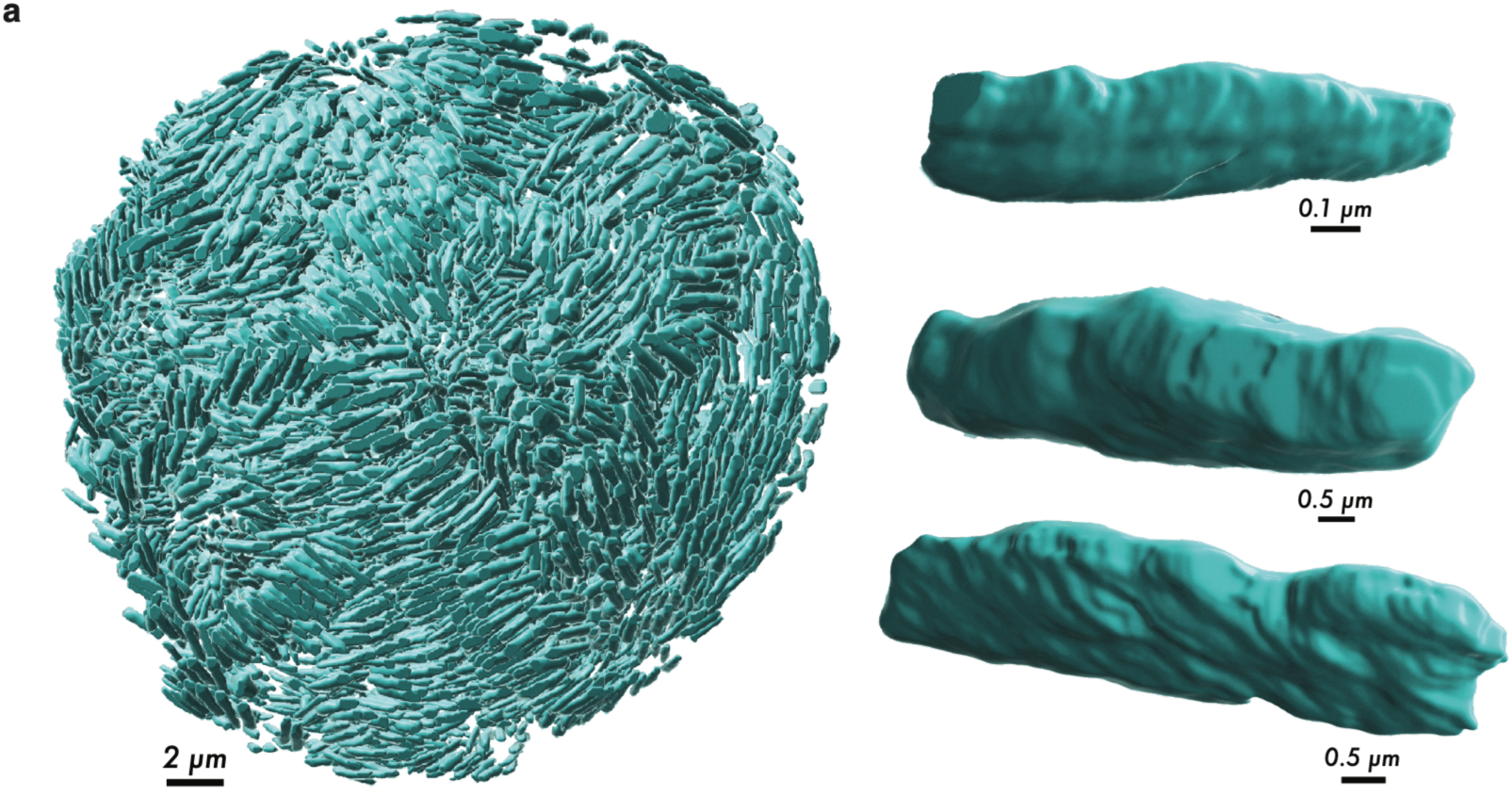
Nuclei in late segmentation are largely uniform in shape and size. **a,** Karyokinesis is complete by late segmentation and partitioned nuclei are elongated and occupy approximately a third of the sporozoite length (all nuclei, left; zoom of subset of nuclei, right).

**Extended Data 3:**
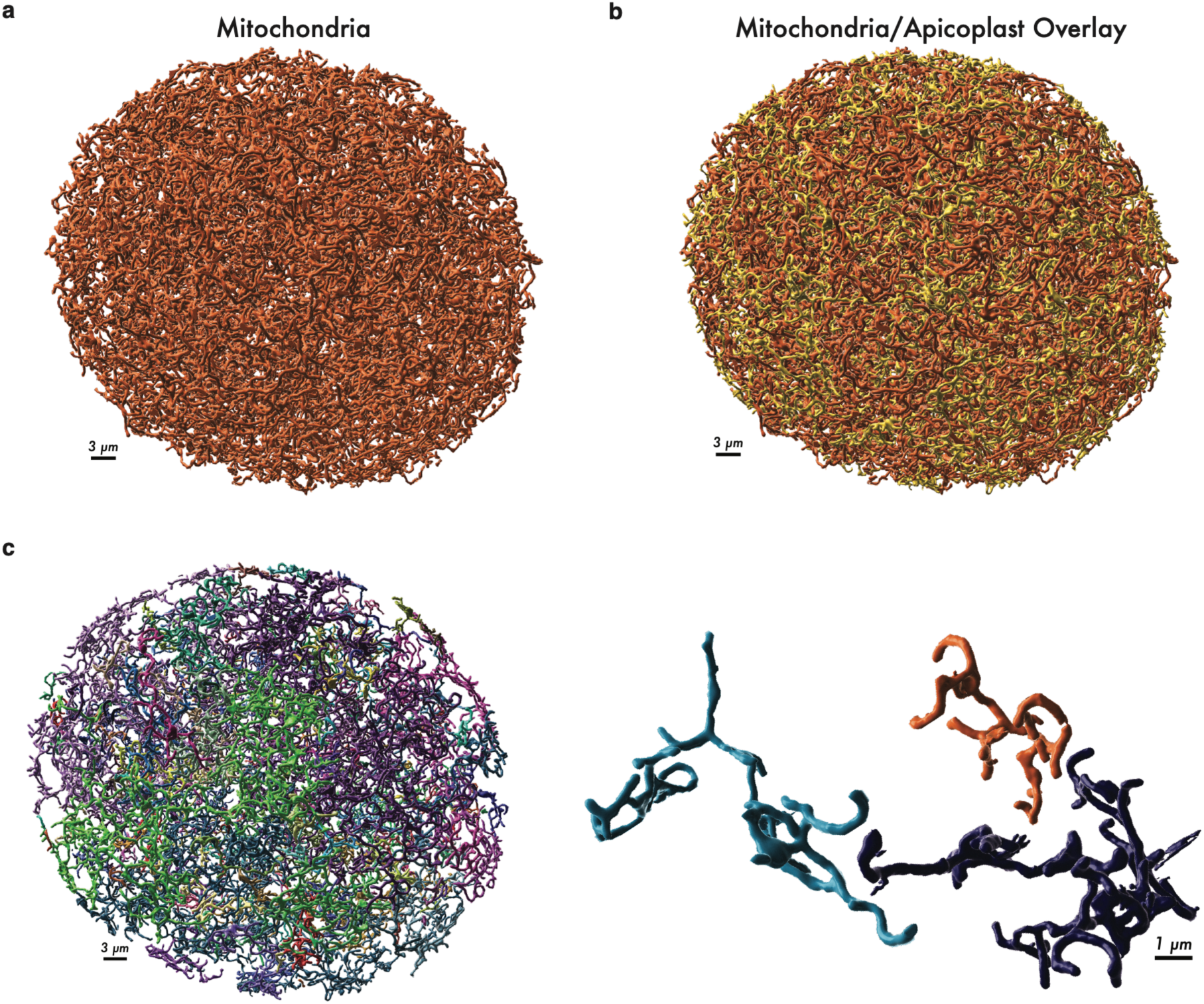
Mid-segmentation mitochondrial and apicoplast morphologies. **a-b,** Mitochondria (*orange)* in mid-segmentation form a convoluted and branching structure that stretches throughout the oocyst volume, similar to the apicoplasts (*yellow*). **c,** Apicoplasts (*color-coded by unique organelle ID*; all plastids, left) in mid-segmentation have already undergone intermediate scission events and 262 discrete apicoplasts were observed. The discontinuity of the apicoplasts is best appreciated when examining a smaller subset of plastids (zoom of subset, right).

**Extended Data 4:**
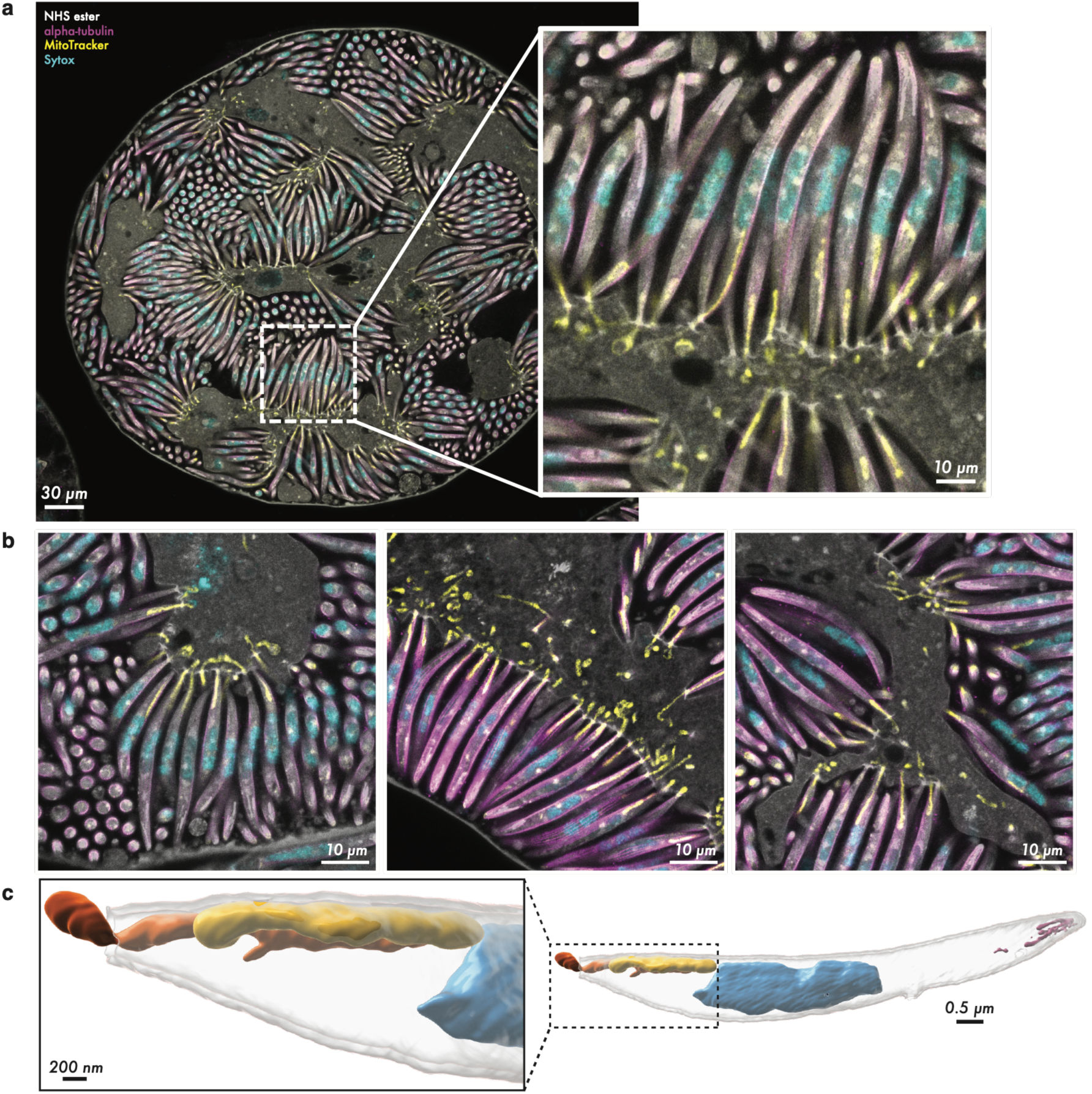
Mitochondria remain tethered to the sporoblast in late segmentation. **a,b** In late segmentation when sporozoites are almost fully elongated (but still attached to the sporoblast at their basal end), the mitochondria can be observed by ultrastructure expansion microscopy as tubular organelles that have yet to undergo their final scission event, protruding past the end of sporozoites into the sporoblast as shown in the inset (**b**). Each inset represents a unique oocyst; all oocysts were analyzed at day 10 post infection. Mitochondria (*yellow*) are stained by MitoTracker Orange CMTMRos (M7510, Invitrogen). The boundaries of the sporozoite and sporoblast are stained by the general protein stain (*grey scale*) AlexaFluor 405 NHS ester (A30000, Invitrogen) while the boundaries of just the sporozoite are highlighted by the presence of alpha-tubulin (*magenta*) (alpha Tubulin monoclonal B-5-1-2; 32-2500, Invitrogen). Nuclei (*cyan*) are stained by SYTOX Deep Red (S11381, Invitrogen). **c,** Mitochondria (*orange*) can be seen extending past end of IMC (*transparent*) reconstructed from FIB-SEM data alongside the apicoplast (*yellow)*, nucleus (*blue*), and rhoptries (*purple*).

**Extended Data 5:**
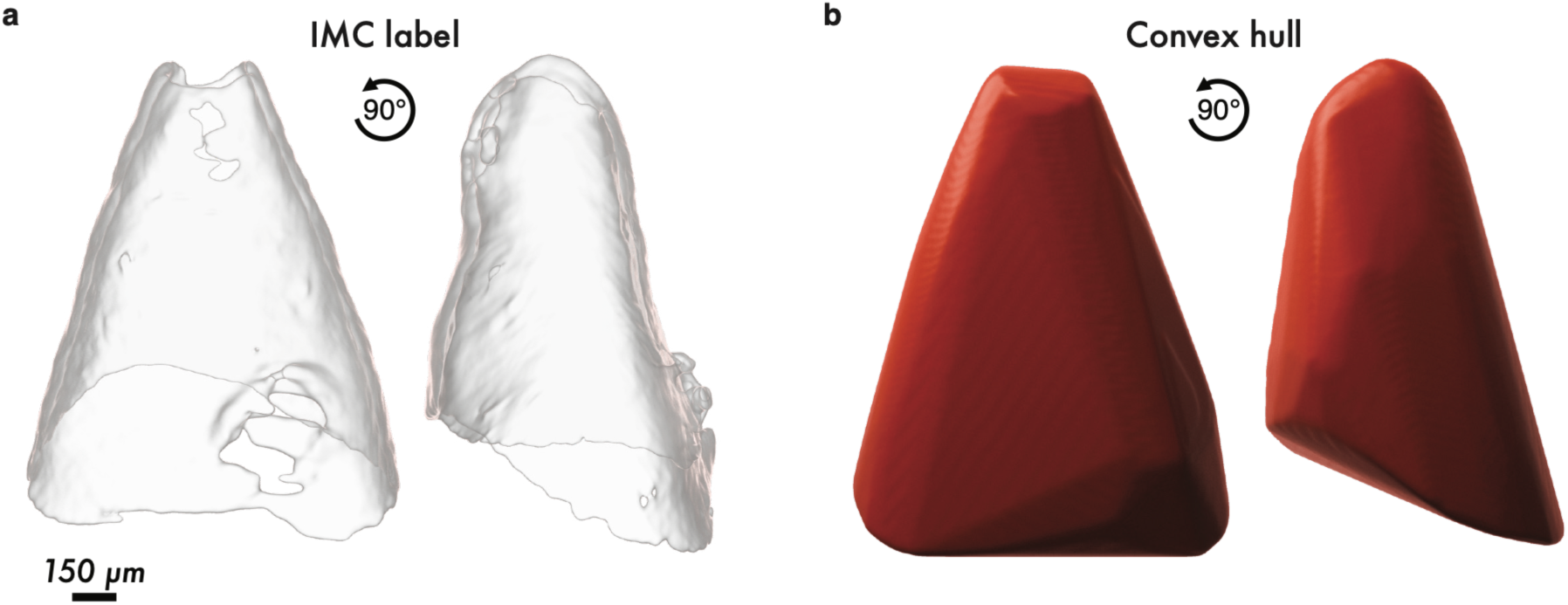
Convex hull accurately models a closed shape for the IMC. **a,** The inner membrane complex (IMC) at mid-segmentation forms an open cone-like shape with a small opening in the top where the apical polar rings resides and a completely open basal end attached to the sporoblast. An example IMC label is shown (*translucent*). The open basal end provides a challenge when quantifying what organelles are contained within the boundaries of the IMC. **b,** Convex hull (*red*) computed in MATLAB (see Methods) closely approximates the IMC label and provides an approximation of the basal boundary.

**Extended Data 6:** Segregation of inherited organelles is imprecise.

|  | 0 | 1 | 2 | ≥3 |
|---|---|---|---|---|
| Mitochondria | 61 | 5,557 | 2,287 | 883 |
| Apicoplast | 77 | 8,588 | 118 | 5 |
| Nuclei | 446 | 8,342 | 0 | 0 |

**Extended Data 7:**
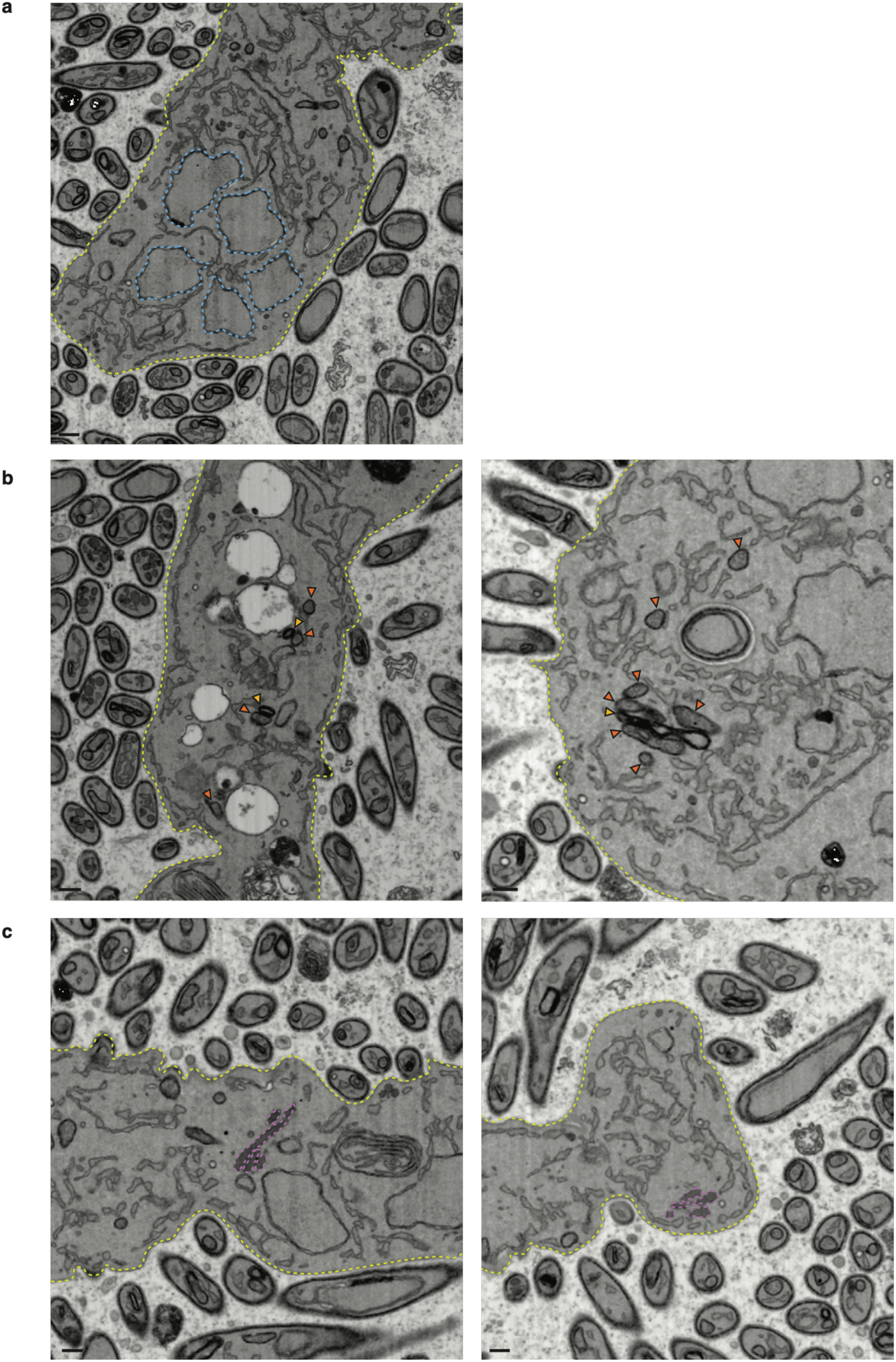
Organelles are retained in the sporoblast during late segmentation. **a,** Residual nuclei (*blue outline*) retained in the sporoblast (*green outline*) appear to be larger in volume than nuclei segregated into daughter sporozoites. **b,** Residual mitochondria (*orange arrowheads*) and apicoplasts (*yellow arrowheads*). Mitochondria appear to be the most abundant residual organelle type by number which may correlate to the greater range in number of mitochondria observed per sporozoite (see Extended Data 7). **c,** Residual rhoptries (*purple outline*) are retained in the sporoblast even when elongation of daughter cells is almost complete. Visually, rhoptries appear to be the least abundant residual organelle type in the sporoblast. Scale bar = 0.2 µm.

