## Supplemental Information Guide for "Nanoscale architecture of malaria sporozoite formation"

**Supplemental Videos**

**Supplemental Video 1:** 3D reconstruction of all organelles in mid-segmentation

In mid-segmentation the apicoplast (*yellow*) and mitochondria (*orange*) form branching, intertwined structures. Immature rhoptries (*purple*) can be seen at the apical ends of newly forming daughter cells and the nuclei (*blue*) are in the process of segregation and the nuclear MTOCs (*dark blue*) are easily visualized. 3D renderings generated from proofread Panoptic DeepLab labels, rendered in Imaris.

**Supplemental Video 2:** 3D reconstruction of all organelles during late segmentation

In late segmentation the apicoplast (*yellow*) is fully segmented and appears as a tubular organelle alongside the mitochondria (*orange*) which has gone intermediate scission steps and similarly appears as an elongated tubular organelle but has not yet undergone the final scission event and remains tethered to the sporoblast. Mature rhoptries (*purple*) are elongated and nuclei (*blue*) are fully segregated and appear more elongated than mid-segmentation. The MTOC is no longer visible at this stage. 3D renderings generated from proofread Panoptic DeepLab labels, rendered in Imaris.

**Supplemental Video 3:** Mid-segmentation FIB-SEM data overview

Overview of raw FIB-SEM data through the entire oocyst volume at mid-segmentation. 3D reconstruction of all IMCs (*white*), rendered in Imaris.

**Supplemental Video 4:** Late segmentation FIB-SEM data overview

Overview of raw FIB-SEM data through the entire oocyst volume at late segmentation. 3D reconstruction of all IMCs (*white*), nuclei (*blue*), rhoptries (*purple*), apicoplast (*yellow*), and mitochondria (*orange*), rendered in MATLAB.

**Supplemental Video 5:** Shifting IMC and organelle morphology during segmentation

For an individual daughter cell, in mid-segmentation the IMC (*transparent*) is cone-shaped, the rhoptries (*purple*) are immature, club-shaped organelles. The nucleus (*blue*) is segregating with the portion containing the nuclear MTOC (*dark blue*) already within the IMC. The apicoplast (*yellow*) is partially contained within the IMC and makes direct contact with the MTOC. The mitochondrion (*orange*) is not found within the IMC and remains in the sporoblast. By late segmentation, the IMC is fully elongated, segregation of the nucleus and apicoplast are complete. The mitochondria have undergone intermediate fission steps and the rhoptries appear mature and elongated.

**Supplemental Video 6:** Mitochondria remained tethered to the sporoblast in late segmentation

The mitochondria (*orange*) has undergone intermediate scission events and appears alongside the apicoplast (*yellow*) as a tubular organelle. Unlike the apicoplast which is fully segregated and contained completely within the boundaries of the IMC (*transparen*t), the mitochondria have not undergone the final scission event and extend past the basal boundary of the IMC into the sporoblast (sporoblast shown in raw EM).

**Supplemental Video 7:** Maturing sporozoites excrete membrane-bound compartments

Late segmentation sporozoites excrete membrane bound compartments from breaks in the IMC (show in raw EM). A protrusion bound by the parasite plasma membrane (*transparent*) can be observed, excreted along the convex curve of the sporozoite and contains a fragment of the mitochondria (*orange*). Parasite plasma membrane was manually labeled, all 3D reconstructions were done using Imaris.
